# Artificial intelligence-enabled automated analysis of transmission electron micrographs to evaluate chemotherapy impact on mitochondrial morphology in triple negative breast cancer

**DOI:** 10.1101/2025.02.19.635300

**Authors:** Argenis Arriojas, Lily M. Baek, Mariah J. Berner, Jiaqi Wang, Joseph Duraisingh, Alexander Zhurkevich, Antentor Othrell Hinton, Matthew D. Meyer, Lacey E. Dobrolecki, Michael T. Lewis, Kourosh Zarringhalam, Gloria V. Echeverria

## Abstract

Advancements in transmission electron microscopy (TEM) have enabled in-depth studies of biological specimens, offering new avenues to large-scale imaging experiments with subcellular resolution. Mitochondrial structure is of growing interest in cancer biology due to its crucial role in regulating the multi-faceted functions of mitochondria. We and others have established the crucial role of mitochondria in triple-negative breast cancer (TNBC), an aggressive subtype of breast cancer with limited therapeutic options. Building upon our previous work demonstrating the functional role of mitochondrial structure dynamics in the metabolic adaptations and survival of chemotherapy-refractory TNBC cells, we sought to extend those findings to a large-scale analysis of transmission electron micrographs. Here we present a novel U-Net artificial intelligence (AI) model for automatic annotation and assessment of mitochondrial morphology and feature quantification. Our model is trained on 11,039 manually annotated mitochondria across 125 micrographs derived from a variety of orthotopic patient-derived xenograft (PDX) mouse model tumors and adherent cell cultures. The model achieves an F1 score of 0.85 on test micrographs at the pixel level. To validate the ability of our model to detect expected mitochondrial structural changes, we utilized micrographs from mouse primary skeletal muscle cells genetically modified to lack Dynamin-related protein 1 (Drp1). We subjected *in vitro* and *in vivo* TNBC models to conventional chemotherapy treatments commonly used for clinical management of TNBC, including doxorubicin, carboplatin, paclitaxel, and docetaxel (DTX). We found substantial within-sample heterogeneity of mitochondrial structure in both *in vitro* and *in vivo* TNBC models. In four of five PDX models, in vivo treatment with DTX elicited significant alteration in mitochondrial elongation and/or area. We went on to compare mammary tumors and matched lung metastases in a highly metastatic PDX model of TNBC, uncovering significant increase in mitochondrial elongation in metastatic lesions compared to their matched primary mammary tumor. The successful application of our AI model to capture mitochondrial structure marks a step forward in high-throughput analysis of mitochondrial structures, enhancing our understanding of how morphological changes may relate to chemotherapy efficacy and mechanism of action. Our large, manually curated electron micrograph dataset - now publicly available - serves as a unique resource for developing, benchmarking, and applying computational models, while further advancing investigations into mitochondrial morphology and its impact on breast cancer biology. This study provides proof of concept that mitochondrial structural remodeling is an additional layer of cellular reprogramming accompanying therapeutic resistance TNBC that merits further investigation.

## INTRODUCTION

Mitochondria are multifaceted organelles responsible for energy production through oxidative phosphorylation (OXPHOS). This process generates ATP, the cell’s primary energy currency, by utilizing electrons derived from nutrients^1^. Beyond energy production, mitochondria play critical roles in regulating various cellular processes, including apoptosis, cell cycle control, and cell signaling, all of which are dependent on their structural integrity^2–5^.

Mitochondrial structure, regulated by their fusion and fission, is highly dynamic. These organelles are enclosed by a double membrane, with the inner membrane folded into cristae that house the OXPHOS machinery^6^. The balance between mitochondrial fusion and fission, governed by key proteins such as dynamin-related protein 1 (DRP1), optic atrophy 1 (OPA1), and mitofusins (MFNs), is crucial for maintaining mitochondrial function and cellular health^7^. In several diseases including cancer, alterations in mitochondrial structure, including those that favor either elongation or fragmentation, are common, contributing to dynamic metabolic reprogramming that supports tumor growth and survival^8–10^. Alterations in mitochondrial fusion and fission can lead to enhanced proliferation, metastasis, or other hallmarks of cancer^9^. Thus, delineating these alterations is critical for gaining a deeper understanding of cancer biology.

Triple-negative breast cancer (TNBC) presents significant treatment challenges due to the absence of estrogen and progesterone receptors and human epidermal growth factor receptor two (HER2) amplification or overexpression^11^. Although neoadjuvant (a.k.a pre-surgical) chemotherapy (NACT) with PD-1 immune checkpoint blockers is the standard treatment, approximately half of patients do not achieve complete tumor eradication, leading to significantly elevated risk of metastasis and mortality^12–14^. In our previous studies, we found that after chemotherapy, residual patient-derived xenograft (PDX) tumors which eventually recurred underwent reversible transcriptional and mitochondrial adaptations, with notable increases in OXPHOS-related gene expression and activity relative to treatment-naive tumors^15^. This mitochondrial adaptation was mediated by elevation of the mitochondrial fusion protein OPA1. We also demonstrated that inhibiting OPA1 with the small molecule MYLS22^16^ abated mitochondrial fusion, OXPHOS, and chemoresistance^8^. Targeting proteins like DRP1, OPA1, or MFNs could enhance the efficacy of existing treatments and overcome resistance mechanisms, offering new hope for patients with difficult-to-treat cancers^17–20^. The role of mitochondrial fission and fusion in TNBC metastasis remains controversial, with contrasting findings reported in differeing experimental models^21–23^.

Electron microscopy is the gold standard method to gain subcellular resolution of organelles. Due to the complexity of mitochondrial structure, accurately quantifying features in imaging data is crucial for understanding the connection between mitochondrial structure and function. Traditionally, analyzing mitochondrial structures involves manually tracing and counting individual mitochondria in electron micrographs to measure morphological parameters such as length, area, circularity, and linearity. This process becomes challenging when the contrast between mitochondria and other organelles or the cytosol is low. In addition, during manual inspection of mitochondria, the examiner often considers only a subset of mitochondria in each image, leading to a loss of information. Individual tumor cells can contain anywhere from a few dozen to several hundred mitochondria, making this manual procedure ineffective for statistically powered high-throughput analysis, as it is extremely time-consuming and may be heavily dependent on examiner bias. These issues can lead to unreliable and inconsistent metrics for quantification, especially considering the extensive intra-tumoral heterogeneity characteristic of most cancers.

In recent years, advancements in artificial intelligence (AI), particularly deep machine learning, have transformed image analysis across various medical domains^24–26^, including those focused on mitochondrial inner and outer membrane structure^27^. Deep learning models, especially convolutional neural networks, have shown significant success in tasks such as tumor classification, cancer subtype identification, mutation prediction, and treatment outcome prediction using histological images like hematoxylin and eosin (H&E) stained slides^28–30^. These approaches utilize either transfer learning, where pre-trained models are fine-tuned for specific tasks, or full-training strategies that train models from scratch using large domain-specific datasets. Such innovations have enabled the identification of subtle morphological and molecular patterns within images that were previously unnoticed through traditional methods. Similarly, AI-driven approaches have advanced mitochondrial structure studies, enabling precise quantification of mitochondrial dynamics and revealing intricate features like cristae topologies that were difficult to detect. Additionally, pixel-level segmentation of cancer images using autoencoder-based architectures has been adapted for high-resolution segmentation tasks across various image types^31–33^. Among these, U-Net architectures^34^ have demonstrated particular efficacy in pixel-level cancer image analysis^28–30,35^. These methods offer automated segmentation and detection capabilities, facilitating more accurate, efficient, and unbiased quantification with exceptional statistical power. Most of these tools are designed for broad organelle segmentation or 3D volume analysis, typically using data from non-cancer tissues. However, an automated tool that could robustly detect mitochondria in TNBC specimens, which are structurally complex with extremely high intra-tumor heterogeneity, are lacking.

In this study, we utilized *in vitro* (adherent cultures of human TNBC cell lines) and *in vivo* (orthotopic PDX mouse models of TNBC) models, in which we previously delineated mitochondrial features and chemotherapeutic responses^15^, to conduct a large-scale assessment of mitochondrial morphology. Building on our studies that demonstrated a link between mitochondrial morphology, OXPHOS, and survival in post-chemotherapy residual TNBC cells^8,15^, we sought to develop and validate an AI-based approach capable of high-throughput segmentation and quantitative analysis of mitochondrial structures in TNBC. To this end, we first generated a large set of transmission electron micrographs from our models, then curated a high-quality dataset by manually annotating 11,039 mitochondria across 125 micrographs from the aforementioned models. These manual annotations served as training data for a U-Net image segmentation model, which was then applied to detect mitochondria at scale. Using computer-vision approaches, we systematically characterized multiple mitochondrial morphological features. This extensive analysis allowed us to quantify mitochondrial structural changes following chemotherapy treatment with high statistical power. Here, we describe the training and evaluation of our model, demonstrate its robustness and versatility, and present findings that illuminate how mitochondrial structure adapts in TNBC. This study establishes that, in addition to transcriptomic and proteomic rewiring long-documented in the literature, chemotherapy actively reshapes mitochondrial morphology, often heterogeneously, across TNBC tumors^15,36–39^. These results identify mitochondrial structural remodeling as a fundamental and previously under-recognized aspect of therapeutic adaptation in TNBC.

## RESULTS

### Diverse training datasets to enable robust mitochondrial segmentation

Using our previously established methods to examine residual cells surviving exposure to conventional chemotherapies with *in vivo* and *in vitro* models of TNBC, we systematically collected cell and tumor specimens then subjected them to TEM. We then assembled a diverse training dataset by manually annotating mitochondria from 125 electron micrographs using QuPath^40^. These images ranged in calibrated pixel size from 0.36 to 23 nm/pixel and were collected from two cell culture models – primary mouse skeletal myotubes genetically engineered to lack the mitochondrial fission-driving gene *Drp1* (*Drp1* KO) and human TNBC cells (MDA-MB-231) as well as seven orthotopic patient-derived xenograft (PDX) TNBC models (PIM001-P, PIM005^15^, PIM056^41^, WHIM-14^42^, BCM-2147^43^, BCM-15116 (unpublished), and HCI-010^42^, each derived from a unique TNBC patient. For each condition, one representative tumor from which multiple fields of view (>50 images per condition) were acquired at varying magnifications was analyzed. At an intermediate resolution of 7.5 nm/pixel, slides typically contained between 100 - 500 mitochondria. Data acquisition involved three separate microscopists, each employing unique microscope settings and capturing various specimens. This diversity in magnification levels, microscope configurations, image formats (including presence of scale bars), and biological systems.

### Model development overview

Figure 1 shows a schematic of our pipeline. Briefly, we began with preparation of training data by manually annotated micrographs, followed by tiling at a common resolution (7.5 nm/pixel), then data augmentation to enhance model robustness. A U-Net was trained, allowing the model to learn complex features associated with mitochondrial morphology. The trained model was then utilized to make predictions on new images, detecting and outlining mitochondrial structures. Finally, we performed downstream analyses in two major steps: (1) computer vision tasks (*e.g.,* gaussian smoothing, contour detection, feature characterization such as maximum Feret’s diameter and area) and (2) statistical assessments (*e.g*., distribution measurements) to compare control versus experimental conditions. No manual proofreading or post hoc correction of segmentation outputs were performed after inference and smoothing. All quantitative analyses were conducted directly on smoothed model-generated outputs. This streamlined workflow laid the foundation for high-throughput, quantitative evaluation of mitochondrial morphology.

**Figure 1.**
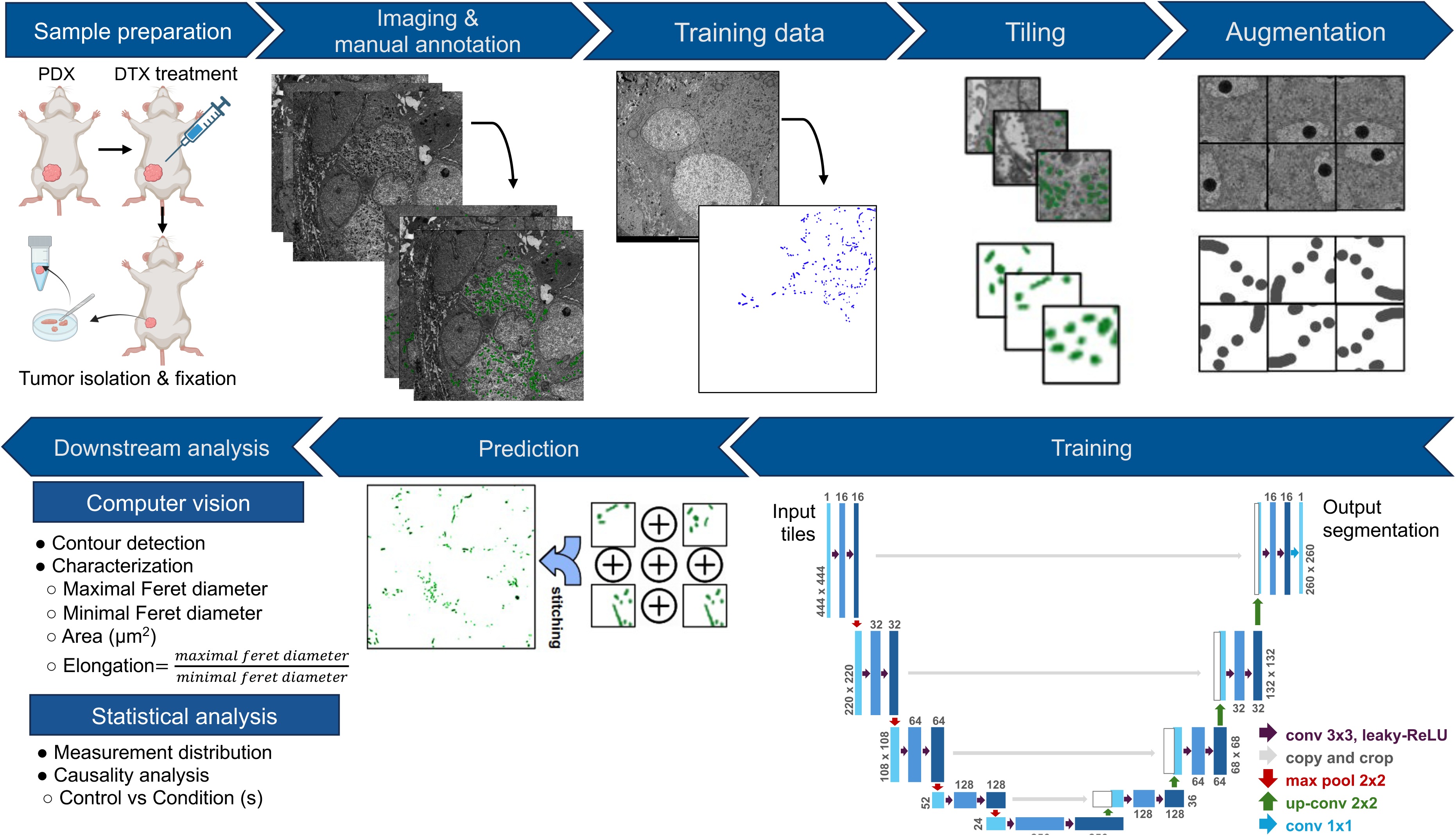
Schematic overview of the U-Net AI pipeline for mitochondrial segmentation and analysis. The workflow begins with the preparation of training data, including transmission electron micrographs and manually annotated masks of mitochondria. Images are then rescaled to a common resolution (7.5nm/pixel) and divided into smaller tiles for input into the neural network. Data augmentation, such as rotations, flips, and brightness adjustments, enhances model robustness to image variability. The U-Net architecture, consisting of contracting and expanding paths, is trained to generate segmentation maps from the input image tiles. Following training, the model is used to make predictions on new images, with segmentation masks stitched together for full-image analysis. Downstream analysis includes computer vision tasks for feature quantification and statistical evaluations of mitochondrial morphology across experimental conditions.

To evaluate the impact of training data composition and model architecture on segmentation performance, we trained and compared several models. Three U-Net models were trained on individual fully annotated datasets (*Drp1* KO, HCI-010, and PIM001-P), each serving as both training and held-out test sets. A fourth U-Net, termed the Mixture model, pooled control images from these three fully annotated datasets along with three additional partially annotated datasets (BCM-2147, MDA-MB-231, and PIM005), totaling 125 micrographs (see Methods). Because the additional datasets in the Mixture were not exhaustively annotated, we evaluated the Mixture model exclusively on the three fully annotated test sets to avoid underestimating performance due to unannotated mitochondria. To explore whether alternative architectures could enhance segmentation, we also trained U-Net++ (Mixture++) and Mask R-CNN (Mixture-RCNN) models using the Mixture training data, and benchmarked all models against the general-purpose MitoNet-v1 segmentation tool (see Methods).

### Model performance evaluation

We employed a 5-fold cross-validation approach, in which each dataset was partitioned into five subsets, with four folds used for training and one fold reserved for validation in an iterative manner (Fig. 2A). This strategy allowed us to evaluate model performance across different splits of the data and assess robustness. As shown in Figure 2B, models were trained independently on each fold and subsequently evaluated on held-out validation data, followed by ensemble averaging across models. Model performance during training was monitored across epochs using F1 score (left panels) and recall (right panels), where F1 score summarizes the balance between precision (the accuracy of predicted mitochondria pixels) and recall (the proportion of true mitochondria pixels correctly identified), while recall specifically reflects the sensitivity of mitochondrial detection. Performance curves across folds demonstrate rapid convergence and consistent behavior across datasets. Importantly, this evaluation reflects robustness within the datasets included in this study and does not imply generalizability beyond these imaging conditions and sample types.

**Figure 2.**
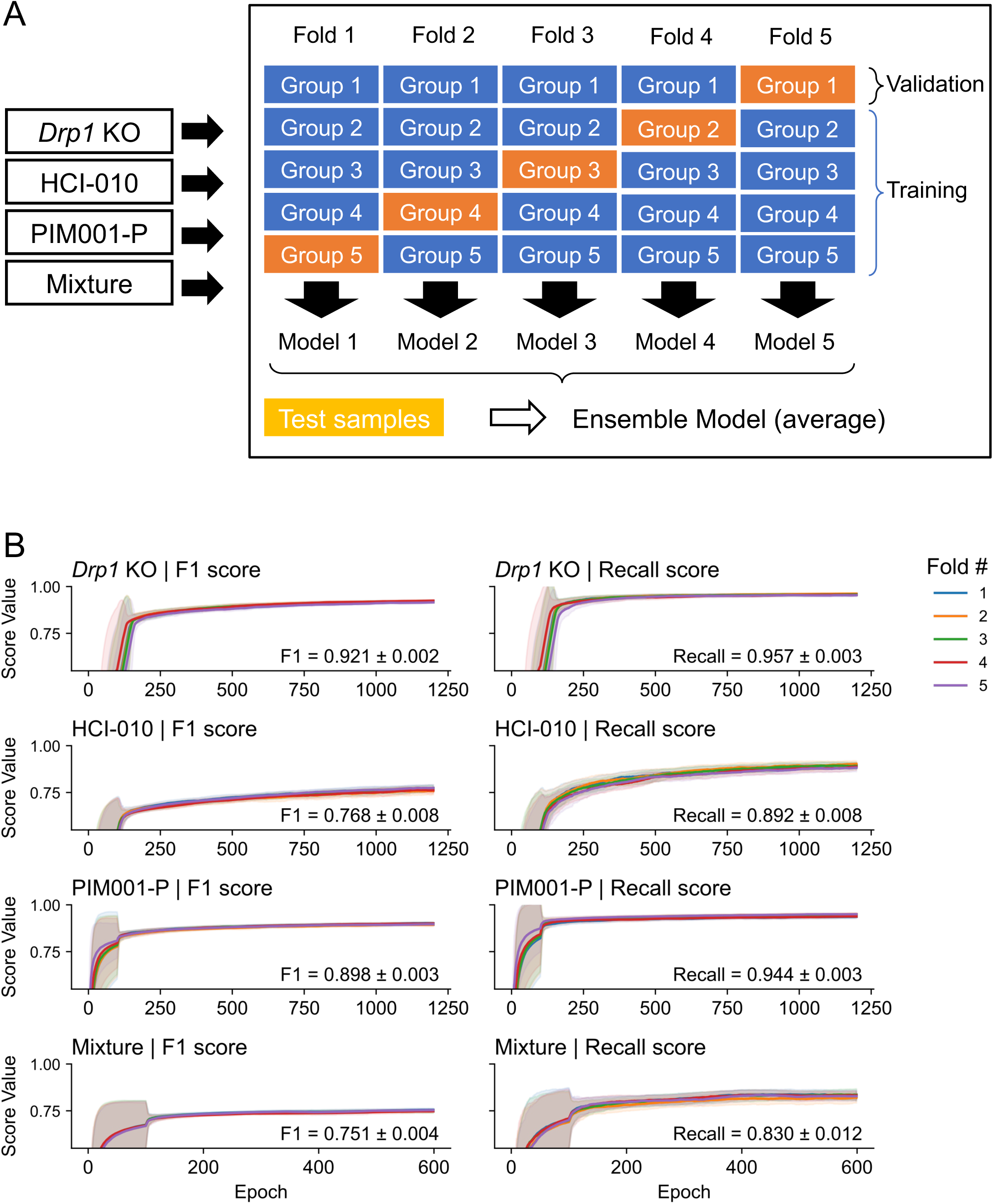
Evaluation of model performance using 5-fold cross-validation across datasets. (A) Schematic of the 5-fold cross-validation strategy. Each dataset (*Drp1* KO, HCI-010, PIM001-P, and a mixed dataset combining samples from all groups) was partitioned into five folds, with four folds used for training (blue rectangles) and one fold reserved for validation (orange rectangles) in each iteration. Models trained on each fold were subsequently combined using an ensemble averaging approach. (B) Model performance during training across epochs. F1 score (left panels) and recall (right panels) are plotted as a function of training epochs for each dataset. Each colored line and shaded region represents the 100-epoch moving average and standard deviation, respectively, for an individual cross-validation fold. F1 score reflects the balance between precision and recall, while recall measures the proportion of true mitochondria pixels correctly identified. Across all datasets, performance rapidly converged and remained consistent across folds, indicating stable model training despite variability in imaging conditions. Together, these results demonstrate consistent model performance across cross-validation splits within each dataset.

We assessed F1 score and recall to evaluate the consistency and robustness of mitochondrial segmentation performance (Fig. 3). The heatmap reveals variations in F1 scores for each dataset, with the PIM001-P PDX model achieving the highest score on its corresponding test dataset (F1 = 0.86), while *Drp1* KO and HCI-010 reached 0.84 and 0.72, respectively, indicating strong agreement between model predictions and manual annotations (Fig. 3A). In contrast, the *Drp1* KO model shows lower performance on other datasets, with scores of 0.53 on HCI-010 and 0.19 on PIM001-P, indicating that dataset-specific characteristics can strongly influence robustness. Training on the Mixture attenuates these differences and yields 0.80 on *Drp1* KO, 0.74 on HCI-010, and 0.84 on PIM001-P, providing competitive performance with improved consistency across datasets. This trend is observed in the evaluation of recall, where the Mixture model performs better than single models trained on individual datasets. While some models achieved higher recall in specific cases, these gains were not consistently accompanied by improvements in F1 score across datasets (Fig. 3A), and qualitative comparisons did not demonstrate a clear advantage in segmentation accuracy (Fig. 3C). Given the class imbalance in this segmentation task, precision–recall (PR) curves are the most appropriate evaluation. The PR AUC values are 0.92, 0.80, and 0.94 for the *Drp1* KO, HCI-010, and PIM001-P models, respectively, which align closely with the F1 scores (Fig. 3B).

**Figure 3.**
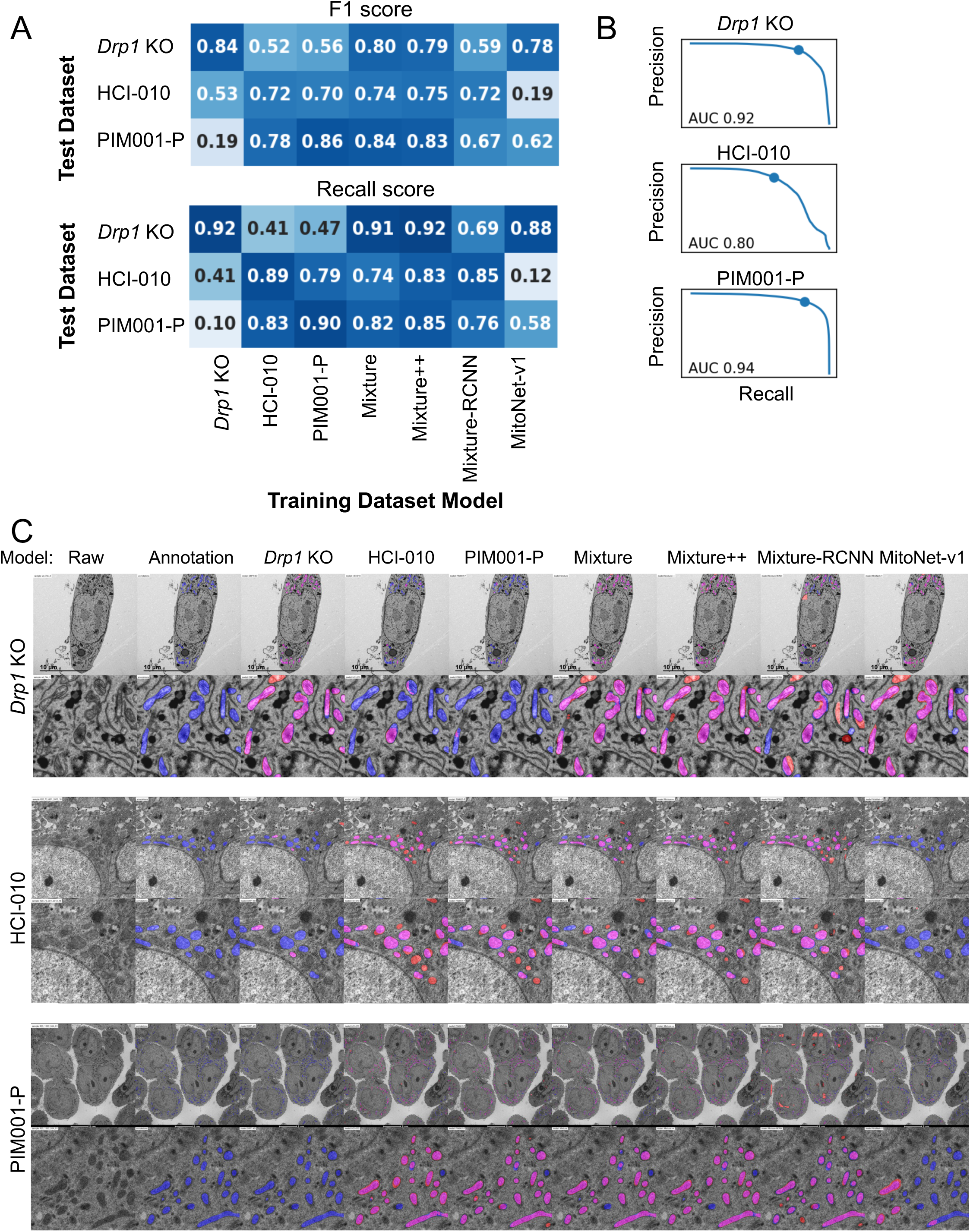
Evaluation of segmentation performance and visual comparison of model predictions. (A) Heatmaps showing F1 scores (left) and recall scores (right) for mitochondrial segmentation models trained on individual datasets (*Drp1* KO, HCI-010, PIM001-P), a pooled dataset (Mixture), or enhanced training strategies (Mixture++, Mixture-RCNN), compared against the benchmark model MitoNet-v1. F1 score reflects the balance between precision and recall, while recall measures the proportion of true mitochondria correctly identified. Each row represents the test dataset, and each column represents the training dataset or model. Higher values (darker blue) indicate better performance. (B) Precision–recall curves for models trained on individual datasets (*Drp1* KO, HCI-010, PIM001-P) when tested on their respective test datasets. Precision (y-axis) is plotted against recall (x-axis) across different decision thresholds. The area under the curve (AUC) is indicated for each model. (C) Representative TEM images showing raw micrographs (left), manual annotations, and mitochondrial segmentation outputs from models trained on individual datasets or mixture-based approaches. Each row represents a different magnification (top: whole cell, middle: subcellular region, bottom: field with multiple cells). Segmented mitochondria are pseudocolored for visualization: manual annotations (blue), model predictions (red), and overlapping regions (magenta). For each magnification, corresponding zoomed-in views of selected regions are provided to better visualize segmentation details and overlap between annotations and model predictions.

To further evaluate predictive accuracy, we visually compared model outputs with manual annotations across datasets. Representative images are shown where mitochondria identified by manual annotation (blue) and model prediction (red) are overlaid (magenta) in the “composed” panels (Fig. 3C). Overall, the model successfully identified the majority of mitochondria, aligning with quantitative results, and demonstrating its ability to reliably detect mitochondrial structure under varied experimental conditions. Despite some challenges in specific datasets, these findings underscore the model’s broad applicability for large-scale mitochondrial morphology analysis, particularly when dealing with mixed image sets.

### Evaluating alternative model architectures

To explore whether other recently developed segmentation architectures could enhance performance, we trained U-Net++^44^ (Mixture++) and Mask R-CNN^45^ (Mixture-RCNN) models using the Mixture training dataset and compared it directly with the corresponding U-Net model. As shown in Fig.3A, U-Net++ achieved similar F1 scores but consistently improved recall across all three datasets, indicating a modest advantage in capturing mitochondria pixels that may be missed by the standard U-Net. In contrast, Mask R-CNN shows substantially lower F1 on all test sets and mixed recall. Although U-Net++ offered modest improvements in recall, these gains were not accompanied by meaningful changes in F1 score. Given that our primary analyses were completed using the U-Net architecture with consistent training, benchmarking, and validation across datasets, we opted to retain the U-Net model for subsequent analyses.

### Model benchmarking against MitoNet

We compared our dataset-specific U-Net models to the general purpose MitoNet-v1 model ^46^ using pixel-level F1 score and recall metrics (Fig. 3A). Across all test datasets, our specialized models outperformed MitoNet-v1, achieving higher F1 scores and recall in most cases. Performance was especially improved for PIM001-P and HCI-010 datasets, where the score gap was the largest. These results demonstrate that domain-specific training on TEM images from TNBC models substantially enhanced segmentation accuracy over a general tool.

### *In vitro* cell models provide biological validations of the algorithm

We sought biological validation of our model with two approaches. First, we genetically manipulated a canonical mitochondrial structure regulator to test the AI’s ability to discern an expected structural difference. Specifically, we conducted TEM of murine skeletal primary cells that were either wild type (WT) or engineered to lack the mitochondrial fission protein Drp1 (*Drp1* KO). Micrographs were analyzed at a resolution of 7.5 nm/pixel. Our data demonstrated a statistically significant increase in elongation (where a value of 1 represents a perfect circle and higher values indicate more elongated shapes) in *Drp1* KO cells compared to WT (Fig. 4B), consistent with the well-established role of Drp1 in causing mitochondrial fission. Histograms (Fig. 4C) show a clear rightward shift in elongation for *Drp1* KO relative to WT, with a broader high-elongation tail, consistent with increased mitochondrial elongation. In contrast, the area distributions are largely overlapping and remain skewed toward small objects, in line with the non-significant difference seen in the Fig. 4B. These findings highlight the ability of our model to effectively capture and quantify expected mitochondrial morphological differences.

**Figure 4.**
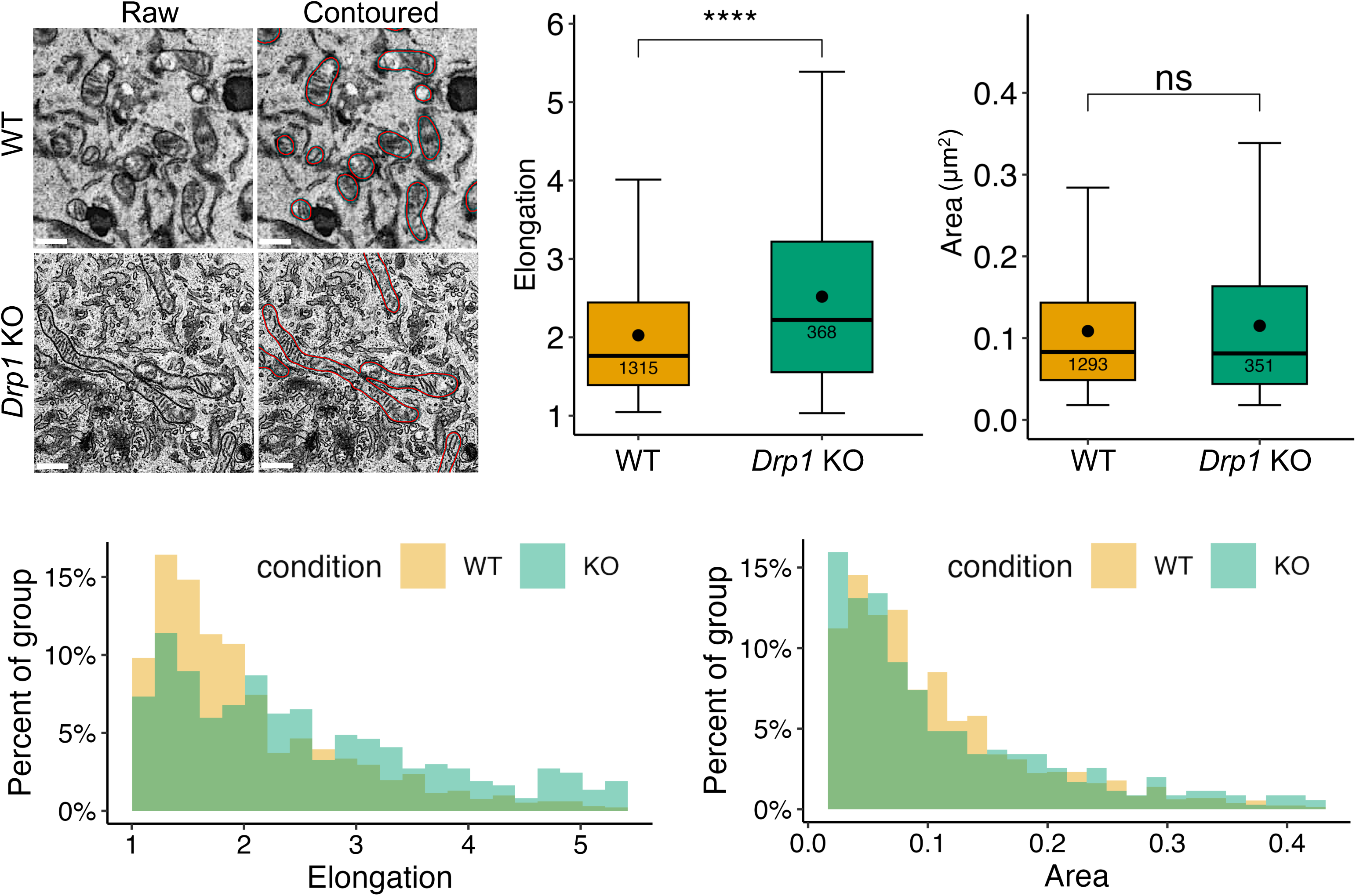
Genetic validation of model performance. (A) Representative micrographs from wild-type (WT) and *Drp1* knockout (KO) murine skeletal muscle cells. Images are shown as raw (left) or with model-predicted mitochondrial contours overlaid in red (right). Scale bars, 0.25 μm. (B) Quantification of mitochondrial elongation (left) and area (µm^2^) (right) in WT and *Drp1* KO cells. Measurements were obtained from segmented mitochondria after converting pixel dimensions to physical units (µm) using image calibration. A total of 42 (WT) and 22 (KO) independent fields of view (FOVs) were analyzed from two biological replicates per condition, with all mitochondria within each FOV included. Boxplots display the mean (black dot), median (horizontal line), interquartile range (box), and whiskers extending to 15–85% anchors, ±1.2×IQR. Mitochondria counts are indicated inside boxes. Data points outside these bounds were excluded as outliers. \*\*\*\**P* < 0.0001, ns= not significant by pairwise Wilcoxon test. (C) Distribution histograms of mitochondrial elongation (left) and area (right) for WT and *Drp1* KO groups, shown as percent of group.

As a second biological validation, we examined micrographs from drug-treated samples for which we had previously established mitochondrial structure alterations using a variety of methods and functional characterizations^8^. We previously reported that different types of standard chemotherapies impacted mitochondrial morphology of TNBC cells in distinct ways: DNA-damaging chemotherapies induced mitochondrial elongation, while taxanes, which disrupt microtubule dynamics, usually reduced mitochondrial elongation (Fig. 5A)^8^. Consistent with those findings, our model detected a significant increase in elongation in MDA-MB-231 TNBC cells treated with DNA-damaging chemotherapies doxorubicin (DXR) or carboplatin (CRB) compared to vehicle (VEH) (Fig. 5B). In contrast, treatment with the taxanes paclitaxel (PTX) or docetaxel (DTX) resulted in decreased mitochondrial elongation and area compared to VEH (Fig.5B). Together, these findings validate our model’s ability to detect expected biological changes and highlight its potential for broader discovery applications.

**Figure 5.**
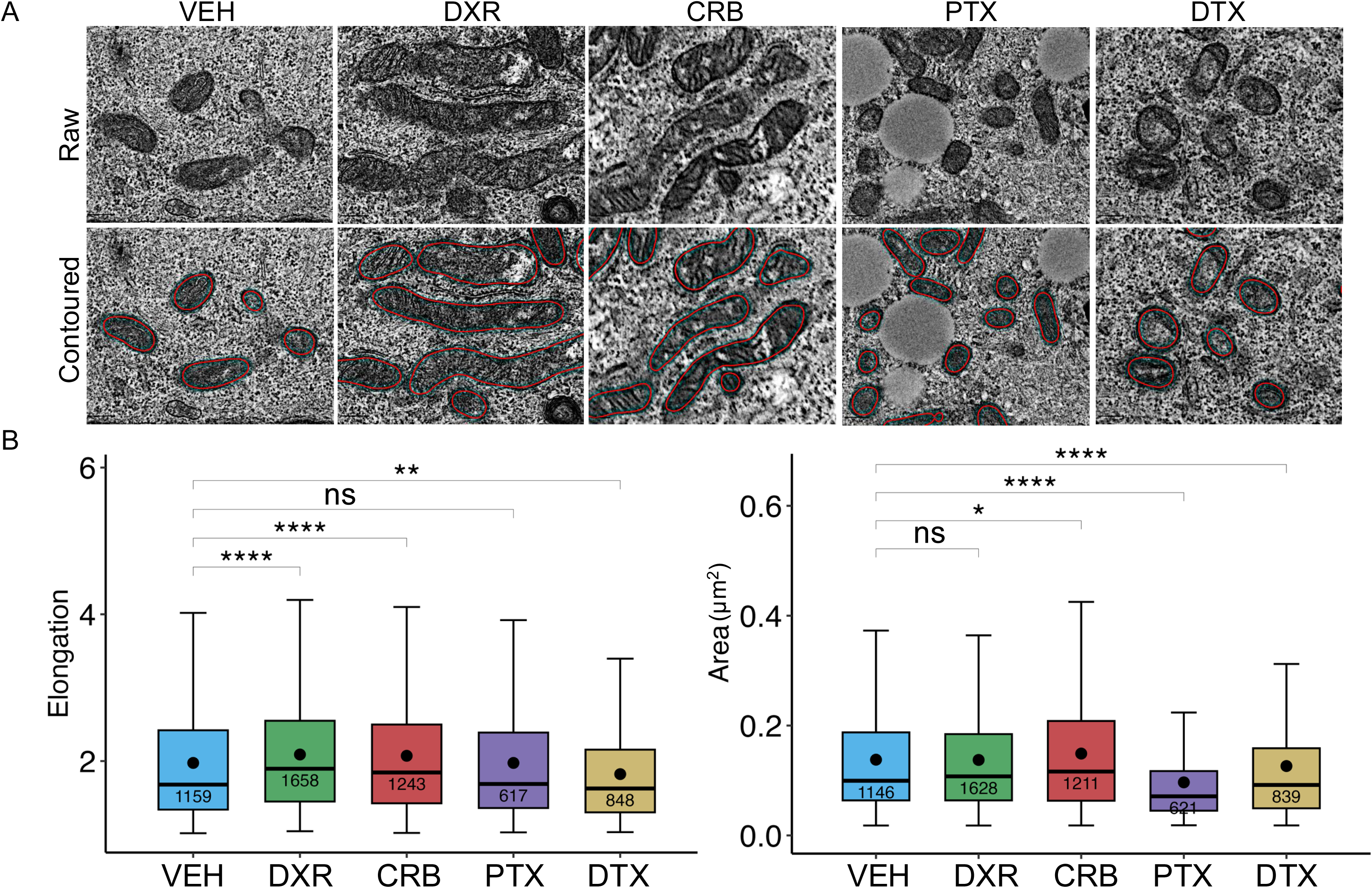
Pharmacologic validation of model performance. (A) Representative micrographs from MDA-MB-231 cells were treated with VEH, DXR, CRB, PTX, or DTX. Images are shown as raw (left) or with model-predicted mitochondrial contours overlaid in red (right). (B) Quantification of mitochondrial elongation (left) and area (µm^2^) (right). Measurements were obtained from segmented mitochondria after converting pixel dimensions to physical units (µm) using image calibration. A total of 85 (VEH), 69 (DXR), 65 (CRB), 44 (PTX), and 82 (DTX) independent fields of view (FOVs) were analyzed from two biological replicates per condition (n = 2), with all mitochondria within each FOV included. Boxplots display the mean (black dot), median (horizontal line), interquartile range (box), and whiskers extending to 15–85% anchors, ±1.2×IQR. Mitochondria counts are indicated inside boxes. Data points outside these bounds were excluded as outliers. \*\*\*\**P* < 0.0001, \*\*\**P* < 0.001, \*\**P* < 0.01, \**P* < 0.05, ns= not significant by pairwise Wilcoxon test.

### Mitochondrial structure changes upon chemotherapy treatment in PDX models

We next sought to investigate whether our approach could detect changes in mitochondrial features, for which we did not have a priori knowledge, during clinically relevant biological transitions in TNBC using orthotopic PDX models. These models are well-known to recapitulate metastatic^47^ and therapeutic response^8,15,36^ phenotypes that match the originating patient tumors. We treated five unique PDX models, representing five TNBC patients, with the conventional taxane agent DTX, often included as a standard treatment for patients with TNBC. Tumors in the five models exhibited varying degrees of responses to DTX (Fig. 6A), as expected based on heterogeneity of chemotherapy efficacy that is common amongst TNBC patients^12^. We collected mammary tumors from mice that were either untreated (n=1-3 per group) or treated with DTX (n=1-3 per group). Residual DTX tumors were harvested 20-28 days after treatment initiation. Tumors were fixed and processed for TEM followed by automated mitochondrial analysis using our algorithm. In three out of five PDX models, we observed significant alteration of mitochondrial elongation following DTX treatment. In four of the five models, significant alteration of mitochondrial area was observed. Representative micrographs illustrate these findings (Fig. 6C). Notably, the magnitude and direction of mitochondrial structural changes varied across models, consistent with the well established inter-patient heterogeneity of TNBC. In models with higher baseline mitochondrial elongation, we observed a tendency toward greater reductions following DTX treatment, although this relationship did not reach statistical significance (Fig. 6D). We did not observe any clear association between mitochondrial features and degree of DTX-induced change in tumor volume (Fig.6A-C). While analyses of these five unique PDX models demonstrate the adaptability of mitochondrial structure, the generalizability of the directionality of those changes to subsets of TNBC patients, perhaps genetically or transcriptomically defined, is yet unclear.

**Figure 6.**
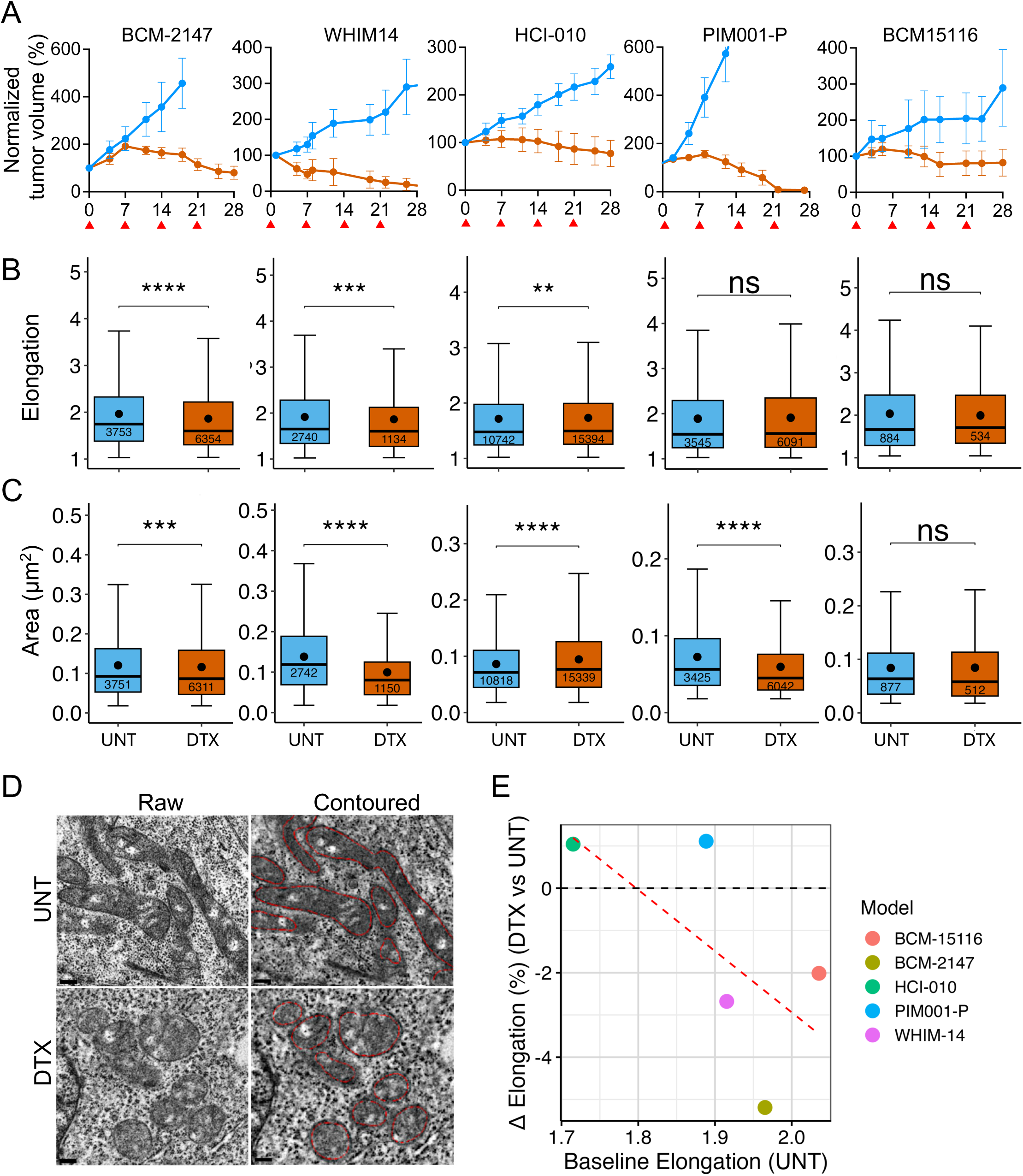
Analysis of mitochondrial changes in PDX models treated with DTX. (A) Line plots showing normalized tumor volume over time in five PDX models treated with DTX (orange) (starting on Day 0) compared to untreated controls (UNT, blue). Red arrows on the x-axis indicate each DTX administration (*i.p*., 20mg/kg) (n≥3). (B-C) Box plots summarize mitochondrial elongation (B) and area (C) across the five PDX models. The number of independent FOVs analyzed per model and condition was as follows: BCM2147, UNT (n = 2, 70 FOVs) and DTX (n = 4, 114 FOVs); HCI-010, UNT (n = 3, 157 FOVs) and DTX (n = 3, 192 FOVs); BCM15116, UNT (n = 1, 52 FOVs) and DTX (n = 1, 53 FOVs); WHIM14, UNT (n = 1, 53 FOVs) and DTX (n = 1, 50 FOVs); and PIM001-P, UNT (n = 1, 66 FOVs) and DTX (n = 1, 89 FOVs). Boxplots display the mean (black dot), median (horizontal line), interquartile range (box), and whiskers extending to 15–85% anchors, ±1.2×IQR. Mitochondria counts are indicated inside boxes. Data points outside these bounds were excluded as outliers. \*\*\*\**P* < 0.0001, \*\*\**P* < 0.001, \*\**P* < 0.01, ns= not significant by pairwise Wilcoxon test. (C) Representative micrographs from BCM-2147 untreated (top) or treated with DTX (bottom). Images are shown as raw (left) or with model-predicted mitochondrial contours overlaid in red (right). Scale bars, 0.2 μm. (E) Scatter plot showing baseline elongation in the untreated (UNT) group (x-axis) vs the percent change (Δ%) in elongation after DTX (y-axis), defined as (DTX_mean_-UNT_mean_)/UNT_mean_ × 100. Each point represents one model. Colors correspond to the legend. The red dashed line indicates the linear regression fit (*P*=0.14), and the black dashed line marks no change.

### Reproducible mitochondrial elongation in a PDX model of spontaneous lung metastasis

We used a highly metastatic PDX model, PIM056, derived from a treatment-naïve TNBC patient. This model develops multifocal macro-metastatic lung lesions within 16 weeks of mammary fat pad (MFP) engraftment without the need for survival tumor resection surgery. This rapidly aggressive metastatic phenotype is exceedingly rare amongst PDX models. Notably, a prior study using H&E and human-specific mitochondrial immunostaining confirmed that the lung metastases in the PIM056 model are predominantly composed of human tumor cells, rather than mouse stromal tissue, thereby validating the cellular identity of the regions analyzed by TEM^48^. Approximately 16 weeks post engraftment, when the mammary tumors measured on average 1000 mm^3^, we collected matched MFP tumors and macro-dissected lung metastases from four individual replicate mice (Fig.7A). TEM followed by automated mitochondria detection revealed significantly longer mitochondria in lung metastases compared to mammary tumors (Fig. 7B-D). In HER2⁺ breast cancer, brain metastases exhibit increased mitochondrial fission relative to MFP tumors^49^. In TNBC, several studies suggest a greater reliance on mitochondrial OXPHOS in lung metastases than at other metastatic sites^50^; however, the contribution of mitochondrial dynamics to these differences remains unresolved. The functional relevance of mitochondrial length in TNBC metastasis will be an important topic for future investigations, and our findings highlight the importance of broadening this analysis to additional models of TNBC metastasis.

**Figure 7.**
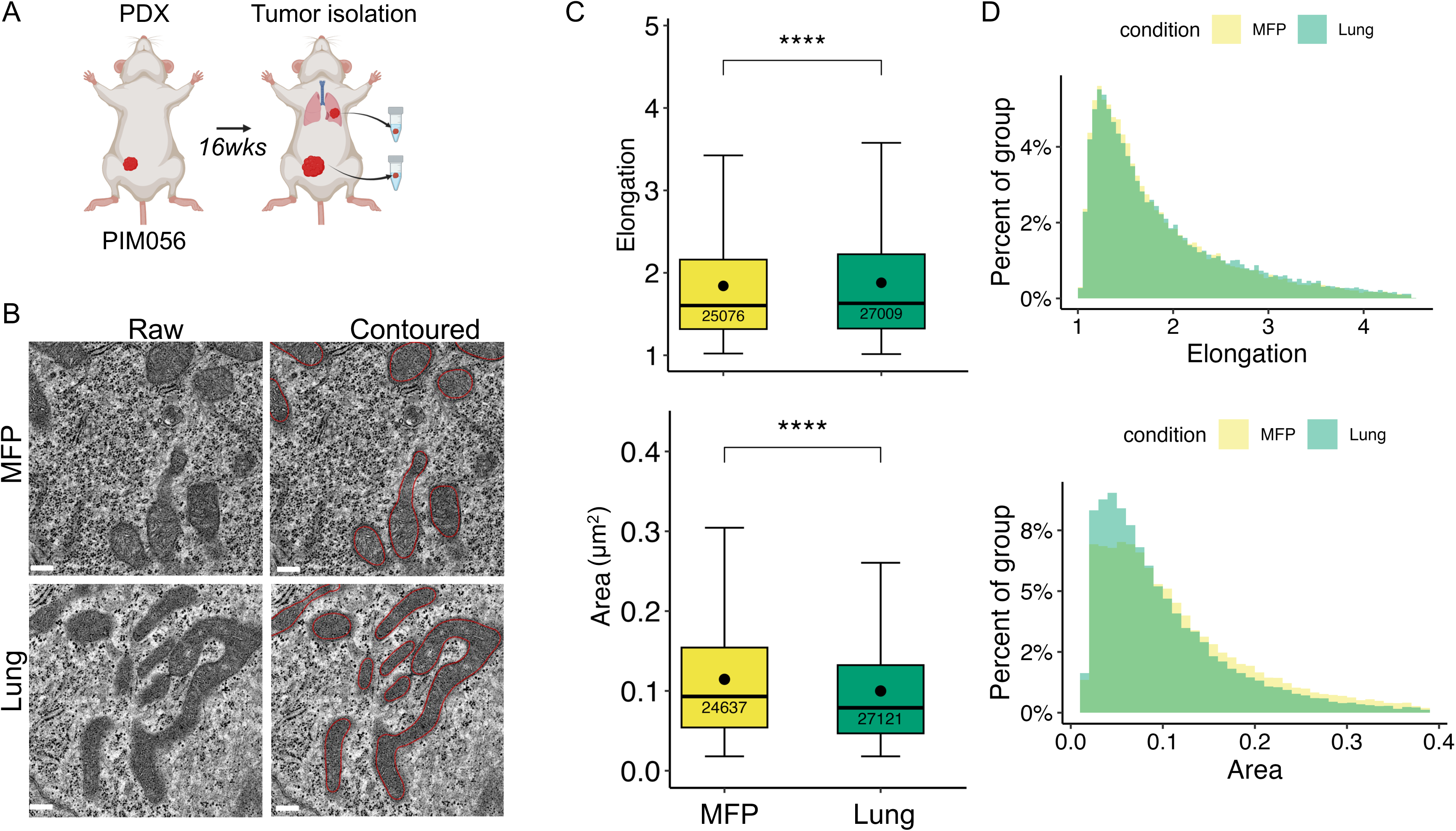
Mitochondrial morphology differences between mammary tumors and lung metastases. (A) Schematic representation of the PIM056 PDX model, where tumors were allowed to grow for 16 weeks before isolation of both primary mammary fat pad (MFP) tumors and lung metastases (Lung) for analysis. At least 60 independent fields of view (FOVs) were analyzed per condition for each biological replicate (n = 4) (B) Representative TEM images show mitochondrial structures in MFP (top) and lung metastases (bottom). Images are shown as raw (left) or with model-predicted mitochondrial contours overlaid in red (right). Scale bars=0.2 μm. (C) Quantification of mitochondrial elongation (top) and area (µm^2^) (bottom) in MFP and lung metastases. Boxplots display the mean (black dot), median (horizontal line), interquartile range (box), and whiskers extending to 15–85% anchors, ±1.2×IQR with mitochondria counts indicated for each group. Data points outside these bounds were excluded as outliers. \*\*\*\**P* < 0.0001 by pairwise Wilcoxon test. (D) Distribution histograms of mitochondrial elongation (top) and area (bottom) for MFP and lung metastases, shown as percent of group.

## DISCUSSION

The application of AI is advancing the field of biomedical image analysis^51^. Manual annotation of cancer images is labor-intensive, prone to human bias, and often inconsistent between analysts, platforms, and institutions, making large-scale studies impractical. The AI-driven model presented here addresses these limitations by automating the detection and characterization of mitochondrial morphology with high accuracy and consistency. A key contribution of our AI model is its capacity to capture and quantify the heterogeneity and dynamics of mitochondrial morphology in biologically important transitions within cancer cells.

Chemotherapy-induced changes in mitochondrial structure are crucial indicators of cellular adaptive mechanisms, including metabolic reprogramming and resistance to cell death. Notably, our prior work demonstrated that genetically or pharmacologically inducing mitochondrial fission or fusion was sufficient to modulate OXPHOS and chemotherapeutic responses^8^. In the present study, we focused on large-scale structural characterization of mitochondrial features and did not directly assess metabolic function. By detecting changes in structural features across large datasets, our approach provides new insights into how mitochondria may contribute to the survival of residual cancer cells post-treatment. Determining how these morphological changes relate to metabolic function, including OXPHOS, will be an important direction for future investigation. We expect our model and highly curated annotations may not only accelerate the analysis of large datasets but may also enable a level of statistical rigor that is difficult to achieve with manual methods. Its robustness, stemming from training on a diverse set of images, underscores its potential for broad applicability in various biological contexts. To benefit the research community, we have made both our algorithm code and our extensive collection of TEM images publicly available. This dataset includes 125 micrographs with a total of 11,039 mitochondrial annotations, which we anticipate will serve as a valuable resource for developing further AI-driven analysis tools.

We observed substantial intra-tumor heterogeneity of mitochondrial shapes and sizes within individual micrographs from both cell lines and PDX models (Supplementary Fig. 1). While this heterogeneity was evident, elongation and area distributions showed substantial overlap across control models, indicating that baseline variability was modest relative to the treatment-associated shifts observed in this study. Although genomic, transcriptomic, epigenomic, and proteomic heterogeneity are well-established phenomena across diverse cancers, including within the cell models in this study, diversity of mitochondrial structure has only recently begun to gain attention in cancer^52,53^. Our algorithm’s ability to capture this heterogeneity with high resolution, even within a decades-old, well-established human cell line MDA-MB-231, suggests an important, previously underappreciated layer of complexity in cancer cell biology. These observations raise compelling questions about how variations in mitochondrial morphology may correlate with functional differences or therapy responses, underscoring the need for deeper investigation into the potential role of mitochondrial heterogeneity in tumor progression.

Our study has several limitations. First, PDX models lack a fully intact immune system, which may be important for mitochondrial features of tumor cells. Consequently, extending our algorithm to immune-competent mouse tumor models will be crucial. Second, the generalizability of our observed chemotherapy-induced mitochondrial structure changes should be further assessed in additional *in vitro* and *in vivo* models with additional conventional chemotherapies, and ideally in human tumor specimens. This is especially important given the well-established inter-patient heterogeneity of TNBC^54–56^. Our algorithm is precisely designed to enable such high-throughput analyses on large sample sets. Third, TEM provides only a two-dimensional cross section of mitochondria and is unable to capture their full three-dimensional complexity^53^. We hypothesize that analyzing high numbers of mitochondria, representing a random sampling of cells’ cross-sections, as presented in this study, may mitigate this limitation by approximating the “average” morphological properties within each sample.

In our metastasis PDX study, we observed altered mitochondrial elongation reproducibly across four replicate mice when comparing their lung metastases to their cognate MFP tumor. These findings are in line with our prior report of the reproducible clonal bottleneck during the process of metastasis in similar TNBC PDX models, and we conducted analogous experiments in biological replicate mice from that model to uncover reproducible alterations in clonal architecture^47^. The fact that mitochondrial changes arose repeatedly in independent mice suggests potential functional relevance, but that must be rigorously tested in future perturbation experiments in PIM056 and in additional models of metastatic TNBC.

We acknowledge that TEM, while providing the gold standard for subcellular resolution of mitochondrial structure, involves intrinsically complex sampling and processing procedures. These include specialized aldehyde fixation, osmium tetroxide post-fixation, resin embedding, ultramicrotomy, and heavy metal counterstaining; procedures that require significantly longer processing times, higher costs, and more advanced technical expertise compared to conventional light microscopy. Despite these practical constraints, TEM remains essential for assessing mitochondrial morphological features, which cannot be resolved with light microscopy. While emerging technologies like focused ion beam scanning electron microscopy (FIB-SEM) enable three-dimensional reconstruction of mitochondrial cristae at nanometer resolution, their application remains even more limited, with single-cell acquisitions requiring days to weeks of imaging and processing^57–59^, restricting throughput to dozens of mitochondria across only a few cells per experiment. This is almost certainly insufficient for capturing the heterogeneity of cancer cell populations. Our AI-driven approach maximizes the value extracted from TEM images by enabling rapid, unbiased analysis of thousands of mitochondria across our manually curated dataset of 125 micrographs, thereby overcoming the traditional bottleneck of manual annotation while maintaining the resolution necessary to detect morphological changes.

While our current analysis focused exclusively on mitochondrial morphology, a more holistic understanding of the chemotherapy-induced cellular changes will require analyzing additional organelles, such as the nucleus, cell membrane, lipid droplets, and endoplasmic reticulum, that can have important interactions with mitochondria. These organelles play significant roles in cellular metabolism, signaling, and homeostasis in concert with mitochondria and may be important contributors to therapeutic resistance in cancer.

Several high-quality datasets and segmentation frameworks have advanced the field of mitochondrial and organelle image analysis, including MitoNet, MitoEM, OpenOrganelle, CREMI, and Lucchi^46,60–64^. Each of these resources serves unique purposes. MitoNet provides a generalizable deep-learning framework for mitochondria segmentation across diverse EM datasets. MitoEM and OpenOrganelle focus on large-scale 3D connectomics data, enabling volumetric reconstruction and detailed mitochondrial network analysis. Meanwhile, the CREMI and Lucchi datasets, though smaller, have served as important benchmarks for EM segmentation methods, often targeting synaptic or nuclear structures alongside mitochondria. Although these tools are invaluable for general benchmarking, they differ substantially from our target application. Most are designed for broad organelle segmentation or 3D volume analysis, typically using isotropic serial-section or FIB-SEM data from non-cancer tissues. By contrast, our pipeline is purpose-built for 2D TEM of TNBC models, with an emphasis on pixel-level morphological quantification of thousands of mitochondria per experiment, enabling the detection of subtle but potentially relevant structural shifts, such as chemotherapy-induced fragmentation or elongation, across large and heterogeneous datasets.

Our rationale for developing a dedicated TNBC-TEM segmentation framework rests on two key points. First, domain-specific optimization was essential, as benchmarking revealed that general-purpose models often underperform on TNBC TEM data due to differences in tissue type, imaging conditions, and structural complexity. Second, our workflow integrates segmentation with downstream morphometric analysis, allowing rigorous statistical comparisons across biological conditions rather than producing masks alone. In this way, our approach complements rather than replaces generalist models, offering a specialized, high-throughput solution for quantitative mitochondrial morphology analysis in clinically relevant cancer models.

We must note that our models were trained exclusively on 2D TEM images from TNBC PDX samples and cell lines and primary mouse skeletal muscle cells, which inherently limits their generalizability. The models are not necessarily expected to produce accurate segmentation predictions for images outside this training domain, such as different cancer types, tissue types, microscopy facilities, or imaging modalities. Therefore, researchers applying our model to new biological contexts should carefully validate the predictions through manual review, and may need to perform additional fine-tuning or retraining with domain-specific annotations to achieve optimal performance.

## METHODS

### TNBC cell culture and drug treatment

The MDA-MB-231 cell line was purchased directly from ATCC (American Type Culture Collection) and cultured in RPMI-1640 (Gibco, 11879020) supplemented with 5 mM glucose (Gibco, A2494001), 10% fetal bovine serum (FBS) (R&D, S11550), and 1× antibiotic-antimycotic solution (Corning, 30-004-CI). Except for carboplatin (solutions were freshly prepared from powder), we prepared stock solutions of 10 mM in DMSO, and further dilutions were prepared in a proper growth medium. Cells were seeded at ∼20% confluence and maintained for 48 h before treatment. Cells were then treated with: doxorubicin (final conc. at 100 nM, Sigma-Aldrich, 44583, protected from light), carboplatin (final conc. at 100 µM, Selleck Chemicals, S1215), paclitaxel (final conc. at 10 nM, Selleck Chemicals, Houston TX, S1150) or docetaxel (final conc. at 5 nM, Selleck Chemicals, Houston TX, S1148) for 48 hours before fixation.

### Authentication of cell lines and PDX models

Cell cultures were tested for mycoplasma contamination each quarter using PCR with the Universal Mycoplasma Detection Kit (ATCC, 30-1012K), and were confirmed to be mycoplasma-free throughout the duration of this study. Short-tandem repeat (STR) DNA fingerprinting was performed on DNA extracted from both cell lines and PDX model tumors by the Cytogenetics and Cell Authentication Core (CCAC) at M.D. Anderson Cancer Center. The Promega 16 High Sensitivity STR Kit (Catalog # DC2100) was used for the fingerprinting analysis, and the resulting profiles were compared against online search databases (DSMZ, ATCC, JCRB, RIKEN) for authentication.

### Animal studies

This study was carried out in accordance with the *Guide for the Care and Use of Laboratory Animals* from the National Institutes of Health (NIH) IACUC. The protocol was approved by the IACUC at BCM (protocol AN-8243 and AN-2289). Mice were euthanized when they reached defined study or ethical end points. Euthanasia was conducted as recommended by the Association for Assessment and Accreditation of Laboratory Animal Care International. PIM001-P, PIM005^15^ and PIM056^41^ were propagated as previously described^1^. Briefly, cryo-preserved PDX cell suspensions were quickly thawed, then washed with Dulbecco’s modified Eagle’s medium (DMEM):F12 (Cytiva HyClone, SH30023.01) supplemented with 5% FBS. Viable cells were counted by staining with AOPI dye (Nexcelom Bioscience, CS2-0106) on a Cellometer K2 (Nexcelom Bioscience). For injection into mammary glands, 0.5-1.0 million viable tumor cells were suspended in a total of 20 µl (1:1 volume mixture of medium and Matrigel (Corning, 354234). Suspensions were then immediately injected unilaterally into the fourth mammary fat pads of 5 to 8-week-old female NOD/SCID [NOD.CB17-Prkdc^scid^/NcrCrl, Charles River, National Cancer Institute (NCI) Colony] or NRG [NOD-Rag1null IL2rgnull] mice (Envigo). For HCI-010, BCM-2147, BCM-15116, and WHIM-14, cryo-preserved tumor fragments (1-2mm^3^) were quickly thawed and washed with DMEM. The tumor chunks were orthotopically xenografted to 4th mammary gland of 6-8-week-old SCID/beige female mice.

Chemotherapy treatment of PDX models was conducted as previously described. Docetaxel solutions were administered by intraperitoneal injection at 20 mg/kg.

For PIM056 metastasis studies, lung metastases and mammary tumors were collected immediately following euthanasia when primary tumors reached approximately 1200 mm^3^. Macro-metastatic lesions visual to the naked eye were isolated with scalpels then preserved for TEM as below.

To generate *Drp1* KO skeletal muscle murine myotubes, mouse husbandry was carried out in accordance with standard protocols^65^ approved by the University of Iowa IACUC. Male C57Bl/6J mice were housed at 22 °C with a 12 h light, 12 h dark cycle and free access to water and standard chow. Controls represent pure C57Bl/6J genetic background mice. In accordance with previous studies^66–68^, Tamoxifen-inducible, skeletal muscle-specific *Drp1* KO mice were generated by crossing mice carrying a homozygous floxed allele of *Drp1* with mice carrying a tamoxifen-inducible Cre recombinase under the control of the skeletal muscle-specific myogenin promoter (Jackson Lab).

### Isolation of satellite cells and differentiation

To collect tissue samples from wild-type or *Drp1* KO mice, animals were anesthetized using isoflurane at 8–10 weeks old. Skeletal muscles from the gastrocnemius and quadriceps were excised and washed twice with 1x phosphate-buffered saline (PBS) supplemented with 1% penicillin-streptomycin and 0.3% fungizone (300 µL/100 mL The muscles were then incubated in Dulbecco’s Modified Eagle’s Medium (DMEM)-F12 containing 0.2% collagenase II (2 mg/mL), 1% penicillin-streptomycin, and 0.3% fungizone (300 µL/100 mL). Samples were shaken for 90 minutes at 37 °C. Following incubation, the media was removed, and the muscle tissue was washed four times with PBS. The media was then replaced with DMEM-F12 containing 0.05% collagenase II (0.5 mg/mL), 1% penicillin-streptomycin, and 0.3% fungizone (300 µL/100 mL), followed by shaking for 30 minutes at 37 °C. Post-digestion, the tissue was ground until cells were dislodged from the tissue matrix and then passed through a 70 µm cell strainer. The isolated cells were centrifuged, resuspended, and plated on BD Matrigel-coated dishes. Adherent cells were differentiated into myotubes by adding DMEM-F12, 20% FBS, 0.004% (40 ng mL-1) basic fibroblast growth factor (R&D Systems, 233-FB/CF), 1× non-essential amino acids, 0.14 × 10^3^ m β-mercaptoethanol, 1x penicillin/streptomycin, and 0.3% fungizone (300 µL/100 mL). Myotubes were maintained in a medium containing 0.001% (10 ng/mL) growth factor until reaching 85% confluency, then were differentiated in DMEM-F12, 2% FBS, and 1× insulin-transferrin-selenium.

### Transmission electron microscopy (TEM)

Tumor cells were fixed, processed, and embedded in situ to prevent morphological damage, as follows. Cells were fixed for 24 hours in Karnovsky’s fixative, post-fixed in 1% osmium tetroxide, dehydrated in graded ethanol, and embedded in EMbed 812 resin. Ultra-thin sections were cut using a Leica UC7 ultramicrotome, stained with saturated methanolic uranyl acetate and Reynold’s lead citrate, and imaged using a JEOL JEM-1230 TEM equipped with an AMT NanoSprint15 sCMOS camera. For tumor tissues, immediately following ethical animal euthanasia, mammary tumors were resected from mice and a ∼1 mm-tick tumor slice was placed in Trump’s fixation solution and fixed for 24 hours at room temperature with rocking. The necrotic areas that usually found in the core region of solid tumors were avoided. The initial fixed samples then were post-fixed in 1% buffered osmium tetroxide, dehydrated in graded ethanol, and embedded in Polybed 812 resin. Ultra-thin sections were obtained using a Leica UC 7 and sections were stained in 1% aqueous uranyl acetate and Reynold’s lead citrate. Images were obtained using a FEI Tecnai Spirit TEM equipped with an Eagle camera and FEI Tia image acquisition software. Approximately 65 images at magnifications of 400-5000x were taken per tumor sample.

Myotubes were processed for TEM in accordance with previous studies^69,70^. For 1 h, cells were fixed by incubating at 37 °C with 2.5% glutaraldehyde in 0.1 m sodium cacodylate buffer. After rinsing twice with 0.1 m sodium cacodylate buffer, samples were fixed again at room temperature for 30 min to 1 h using 1% osmium tetroxide and 1.5% potassium ferrocyanide in 0.1 m sodium cacodylate buffer.

After secondary fixation, samples were washed for 5 min with 0.1 m sodium cacodylate buffer (7.3 pH). From there, two washings of 5 min with diH_2_O ensured the plates were cleaned. While keeping all solutions and plates at room temperature, the samples were incubated with 2.5% uranyl acetate, diluted with H_2_O, at 4 °C overnight. Following this, samples were dehydrated using an ethanol gradient series. After dehydration, the ethanol was replaced with Eponate 12 mixed in 100% ethanol in a 1:1 solution, then incubated at room temperature for 30 min. This was repeated three times for 1 h using 100% Eponate 12. The plates were finally placed in new media and cured in an oven at 70 °C overnight.

Plates were cracked upon hardening, and the cells were separated by submerging the plate in liquid nitrogen. An 80 nm thickness jeweler’s saw was used to cut the block to fit in a Leica UC6 ultramicrotome sample holder. From there, the section was placed on formvar-coated copper grids. These grids were counterstained in 2% uranyl acetate for 2 min. Then these grids were counterstained by Reynold’s lead citrate for 2 min. Images were acquired by TEM on either a JEOL JEM-1230, operating at 120 kV, or a JEOL 1400, operating at 80 kV.

### Manual annotation of micrographs

For each tumor or cell line, 50-60 transmission electron micrographs were obtained at different magnifications, ranging from 1000X to 5000X magnification. These micrographs were then loaded into the QuPath software^40^, a versatile tool for bioimage analysis. Each mitochondrion within the micrographs was manually annotated using the drawing tool provided by QuPath. This process involved carefully outlining the boundaries of individual mitochondria to ensure precise and accurate characterization of their morphological features. The manual annotations were subsequently used for further analysis, providing a detailed dataset for evaluating mitochondrial structure across various tumor samples.

### Model

A U-Net model was utilized to perform semantic segmentation on the TEM images. This deep-learning model is an autoencoder that consists of a feature extraction stage (contracting path, encoder) and a reconstruction stage (expansive path, decoder). The model uses a series of convolutional and max pooling layers to transform the input image into abstract features in a latent space, that will then be decoded in the reconstruction stage to match the model output with the corresponding training mask. A defining characteristic of a U-Net model is the use of feature maps from the contracting path, as input features in the expansive path. This choice helps to localize and assemble a more precise output mask^34^. The network architecture is described in the following. The 2D input layer is followed by a Gaussian Noise layer with standard deviation of 0.01. Each convolutional layer has kernel size 3 and is followed by a leaky ReLu activation layer, a 2D spatial dropout layer (rates 0.2 and 0.1 in the contracting and expansive paths respectively), and a batch normalization layer. A convolutional block is composed of two such convolutional layers, followed by either a 2 by 2 max pooling layer in the contracting path, or a 2 by 2 up-convolution layer with stride 2 in the expanding path. The model is composed of 9 convolutional blocks in total: 4 in the contracting path, 1 in the bottleneck, and 4 in the expansive path. The first convolutional block uses 16 filters in each of its convolutional layers. The number of filters used in the convolutional layers doubles after each max pooling layer, and halves after each up-convolution layer. The full model has just under 2 million trainable parameters.

### Datasets

Three fully annotated datasets were used for training and testing: *Drp1* KO (26 images), HCI-010 (21 images), and PIM001-P (37 images), with the latter two derived from *in vivo* PDX mouse models. All images correspond to non-perturbed (control/vehicle) samples. A U-Net model was trained independently on each dataset. A fourth Mixture model was trained by pooling these three datasets along with partially annotated control images from three additional datasets (BCM-2147, MDA-MB-231, and PIM005), totaling 125 images. Because the additional datasets were not exhaustively annotated, model evaluation was restricted to the three fully annotated test sets.

### Preprocessing training data

The training dataset consists of electron micrographs, each paired with a manually annotated segmentation mask indicating the mitochondria. Magnification information was parsed from the image to obtain the calibrated pixel size. Prior to training or inference, images were scaled to match a common pixel size of 7.5 nm/pixel. Each image-mask pair was tiled to appropriately sized input samples that fit the model architecture shape (444 by 444 in the input layer, and 260 by 260 pixels at the output). The tiled process was performed in an overlapping fashion with a stride of 130 pixels in both horizontal and vertical directions. To avoid losing the pixels at the border, the images were zero-padded. The pixel values are rescaled to lie in the range [0,1]. The tile sets were converted into TF records to improve training computational performance. The final output was assembled by stitching the overlapping predicted mask tiles and taking the average for the overlapping regions.

### Data augmentation

During model training, the input tiles were randomly transformed by rotating and flipping the images. This step was performed to increase the number of training samples and to avoid orientation bias. Moreover, the brightness, contrast and gamma image parameters were randomly adjusted to account for variation of these parameters across the input images. Additionally, images were randomly magnified to boost the robustness of the model to input image resolution.

### Model training

The models were trained on Nvidia DGX-A100 GPUs, for a total of 1200 epochs (600 epochs for the Mixture models) and batch size 128. The Adam optimizer was used with the default parameters, and a weighted binary cross entropy loss function, assigning double weight to positive mask pixels. Model checkpoints were saved for the best achieved loss on the validation sets. The convergence of the model was monitored using the corresponding loss metric, as well as the IOU and DICE coefficients.

### Performance evaluation

The data was split into a training set and a test set (85% and 15% approximately). A 5-fold cross validation was used in the training set, where each fold uses 4 subsets for training and the remaining 5th set as validation. Model performance was assessed using the left-out test set.

### Model predictions

Each of the 5 resulting models from the cross-validation procedure were used to make predictions on new images. An ensemble approach was used to aggregate the results in a single prediction, by taking the average of all predictions. Since the semantic segmentation models are optimized for analysis at a calibrated pixel size of 7.5 nm/pixel, input images are scaled to this magnification prior inference, and predictions are scaled back to the original size.

### Assesing alternative model architectures

Our U-Net implementation was extended to the more recent U-Net++ architecture to evaluate whether its enhanced skip connections could improve segmentation performance. The model was constructed using the same parameters as the standard U-Net, resulting in approximately 2.4 million parameters in total, a ∼25% increase compared to the original architecture.

We also evaluated a general-purpose instance segmentation framework, Detectron2’s COCO-pretrained Mask R-CNN (mask_rcnn_R_50_FPN_3x, Meta AI), to assess whether this architecture could offer improvements over our U-Net-based models. For compatibility, the ground truth annotations from our Mixture training dataset were prepared following Detectron2’s requirements, converted from semantic masks into COCO-style JSON format containing per-instance metadata such as contour coordinates and instance IDs, and registered in Detectron2 as a new local dataset. Model inference was performed via a tile-wise pipeline (512 by 512 px tiles with 20% overlap), followed by stitching to full resolution and rasterizing instance predictions into semantic masks. Despite its versatility, the Mask R-CNN baseline did not outperform our domain-specific U-Net models on the test datasets.

### Benchmarks

To benchmark our model’s performance, we compared it against MitoNet_v1^46^, a widely-used mitochondria segmentation tool. The comparison was performed using the Empanada wrapper (v1.2) integrated with Napari (v0.4.18) in a Python 3.9 conda environment. We applied MitoNet’s 2D inference tool with default parameters to 13 test images from our dataset, generating instance segmentation masks for mitochondria in each image. To enable direct comparison with our U-Net predictions, the instance segmentation masks were converted to semantic segmentation format by flattening individual mitochondria masks using Napari’s “convert to image” feature. Model performance was quantified using pixel-level F1 score and recall metrics.

### Postprocessing for morphometric analysis

For morphological measurements, predicted binary masks were first smoothed to reduce single-pixel noise and produce cleaner object boundaries. This was achieved by repeated convolution with a normalized 2D Gaussian kernel (sigma = 5 nm in image space) for 24 iterations, followed by by re-thresholding to recover a binary mask. This smoothing step was applied uniformly to all predictions prior to morphometric quantification, and was not used in the calculation of segmentation performance metrics.

### Measurements and statistical analysis

For each detected mitochondrion, we compute the cross-sectional area and an elongation score defined as *P*^2^ / *4*π*A* (commonly referred to as a shape factor or inverse circularity), where *P* and *A* are the perimeter and the area of the instance mask. The score equals 1 for a circular shape and increases as the shape departs from circularity. Given that the obtained contours are smooth, this quantity is dominated by elongation rather than boundary irregularity. For each group, data are summarized as the median, mean (black dot in boxplots), interquartile range (IQR), and whiskers spanning the 15th–85th percentiles with ±1.2×IQR, unless otherwise noted. Differences between groups were assessed using the Wilcoxon rank-sum test (two-sided) because the distributions of elongation and area are non-Gaussian, highly skewed, and contain unequal variances and outlier structures. The segmentation pipeline supports both U-Net and U-Net++ architectures; we used U-Net throughout, as the two produced equivalent area and elongation measurements across representative datasets (Table S1).

## Data and code availability

Cell line models of TNBC are available for purchase from ATCC. PDX models can be made available to investigators through a material transfer agreement with the University of Texas MD Anderson Cancer Center (PIM001-P, PIM005 and PIM056) or Baylor College of Medicine (HCI-010, BCM-2147, BCM-15116, and WHIM-14). The Python code is available on GitHub [https://github.com/umbibio/tem-seg] (DOI: 10.5281/zenodo.19442010). Electron micrographs and annotated mitochondrial masks are deposited in Bioimage Archive^71^ under accession # S-BIAD2271. Training datasets and model weights are available on Zenodo with DOIs <u>10.5281/zenodo.15602047</u> and <u>10.5281/zenodo.15602445</u> respectively.

**Clinical trial number**: not applicable

## AUTHOR CONTRIBUTIONS

A.A., L.M.B., K.Z., and G.V.E were responsible for overall completion of the study, writing the manuscript, and preparation of figures.

L.M.B., K.Z., and G.V.E were responsible for project conceptualization.

L.M.B. conducted PDX and cell line experiments, generated electron micrographs, conducted manual annotations of mitochondria, and data analyses under the supervision of G.V.E.

A.A. conducted computational algorithm development, optimization, and testing under the supervision of K.Z.

M.J.B. conducted PDX chemotherapy trials and generated electron micrographs analyzed in this study under the supervision of G.V.E.

A.Z. conducted initial algorithm development under the supervision of K.Z.

J.W. and J.D. contributed to the benchmarking efforts. J.W. performed training and evaluation for the Mixture-RCNN model. J.D. performed model evaluations for the MitoNet-v1 segmentation model.

A.O.H. provided electron micrographs of WT and *Drp1*KO myotubes.

JW implemented the Mask-RCNN model and performed benchmark analysis.

JD implemented MitoNet and Empanada benchmakrs.

M.D.M. aided in design of electron microscopy imaging and generated most images in this study.

L.E.D. conducted PDX chemotherapy trials and provided tissues for electron microscopy under the supervision of M.T.L.

## ACKNOWLEDGEMENTS

We are grateful to the breast cancer patients who donated their biopsies for cell line and PDX model generation. We are grateful to Mrs. Janice Cowden for providing advocacy support for our research. Dr. James P. Barrish assisted with TEM. Dr. Junegoo Lee assisted with mouse experiments. Dr. Helen Piwnica-Worms provided the PIM001-P and PIM056 PDXs through MTA with the University of Texas MD Anderson Cancer Center, and model generation was supported by a generous gift from the Cazalot Foundation through the MD Anderson Women’s Cancer Moonshot Program. STR DNA fingerprinting was done by the Cytogenetics and Cell Authentication Core at M.D. Anderson Cancer Center.

