## Supplementary figures and images for "Artificial intelligence-enabled automated analysis of transmission electron micrographs to evaluate chemotherapy impact on mitochondrial morphology in triple negative breast cancer"

### Supplementary Figure 1

#### Supplementary Figure 1

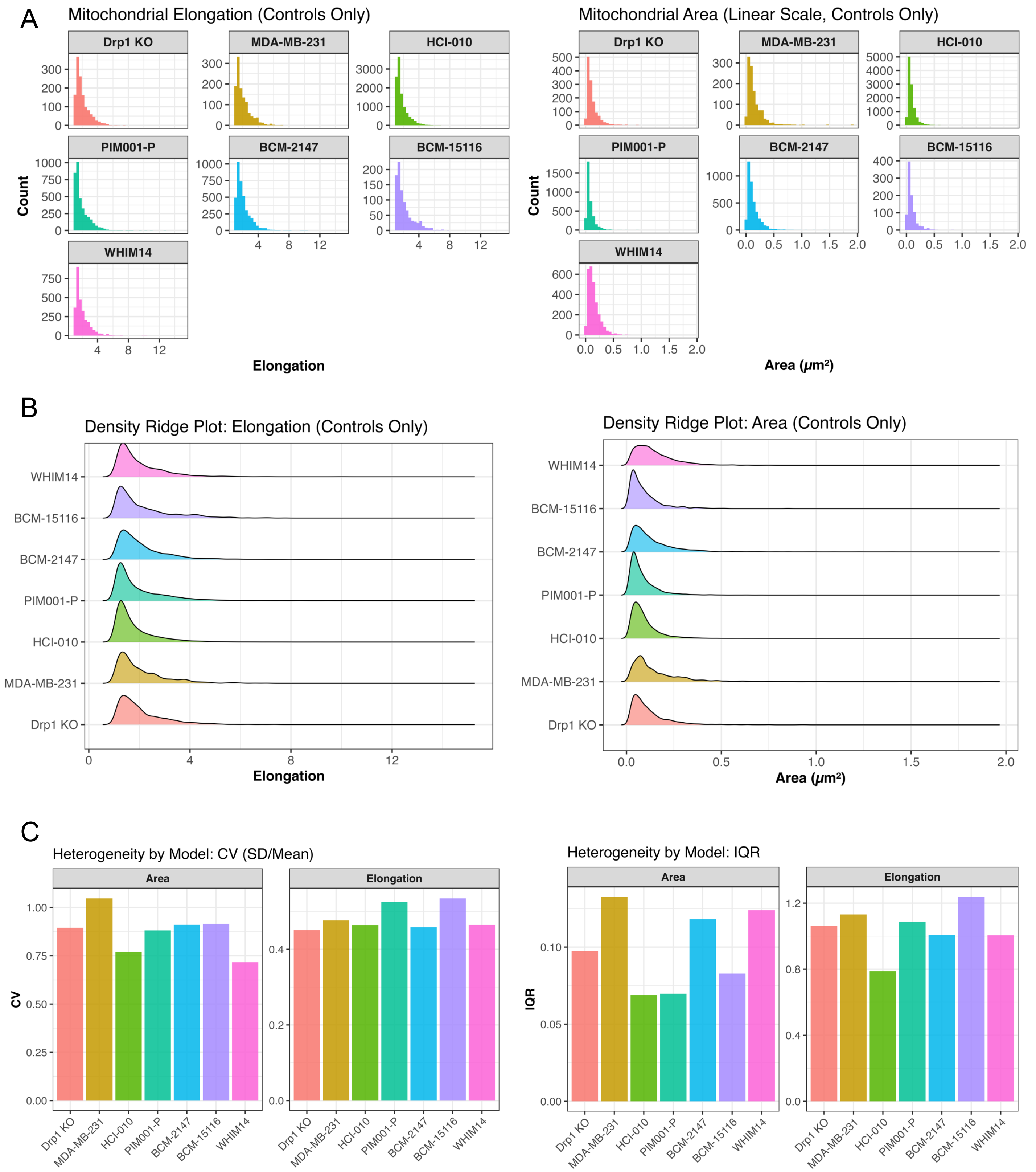

### Supplementary Figure 2

#### Supplementary Figure 2

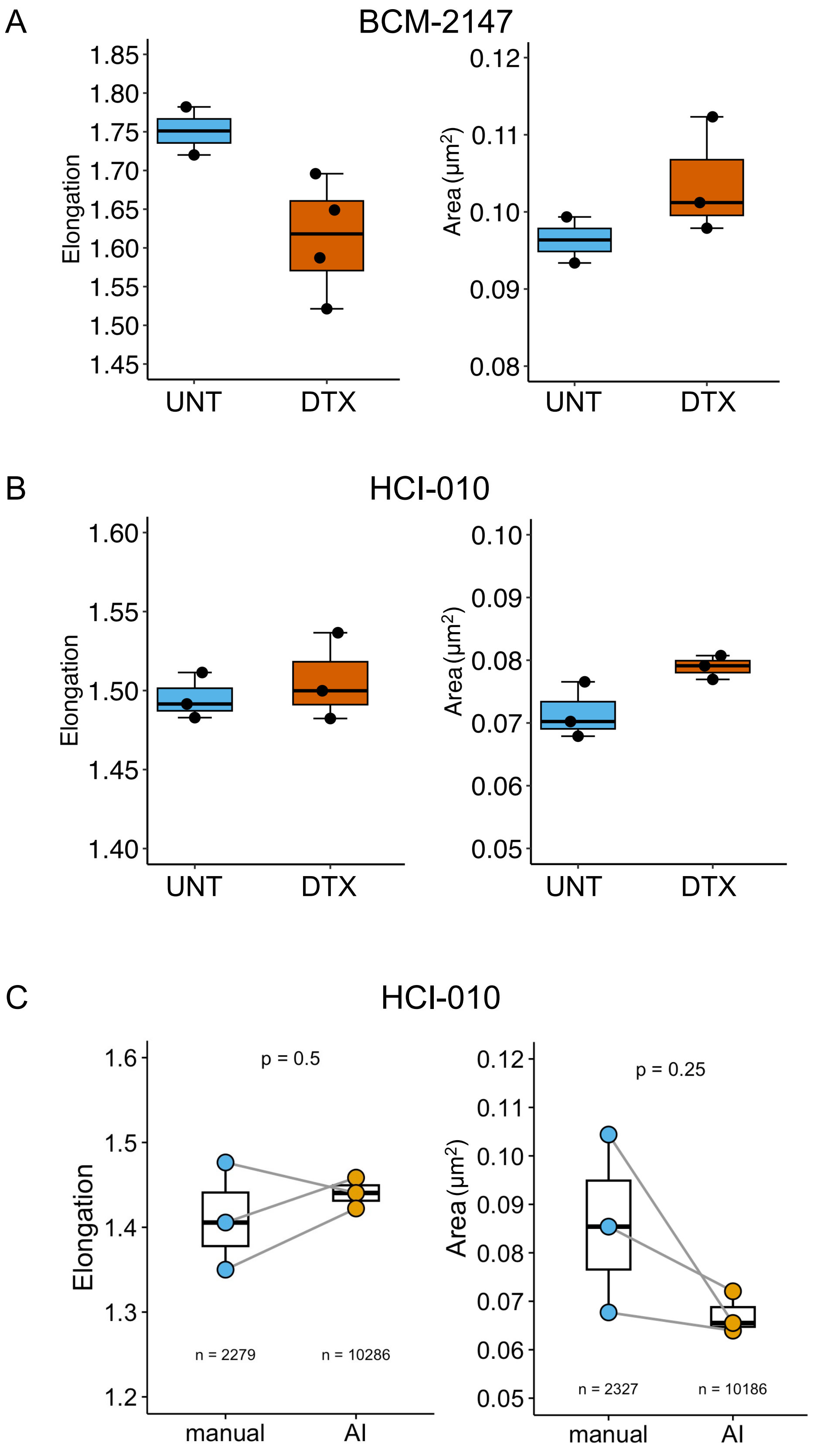
