## Supplementary Table 1 for "Artificial intelligence-enabled automated analysis of transmission electron micrographs to evaluate chemotherapy impact on mitochondrial morphology in triple negative breast cancer"

**Table S1. U-Net and U-Net++ yield equivalent area and elongation measurements and concordant biological conclusions in three representative datasets.**

| **Dataset (comparison)** | **Metric** | **Cross-model agreement (Pearson *r*)** | | **Δ median %**  **U-Net / U-Net++** | **Cliff’s δ**  **U-Net / U-Net++** | **Mann–Whitney *p***  **U-Net / U-Net++** |
| --- | --- | --- | --- | --- | --- | --- |
|  |  | **per-sample *r*** | **paired per-object *r*** |  |  |  |
| HCI-010 (DTX vs. vehicle) | *area* | 0.98 | 0.96 | +12.7 / +8.8 | 0.08 / 0.06 | < 0.001 / < 0.001 |
| HCI-010 (DTX vs. vehicle) | *elongation* | 0.94 | 0.82 | +1.1 / +1.5 | 0.05 / 0.07 | < 0.001 / < 0.001 |
| BCM-2147 (DTX vs. vehicle) | *area* | 0.95 | 0.93 | −5.1 / −3.0 | −0.04 / −0.03 | 0.002 / 0.012 |
| BCM-2147 (DTX vs. vehicle) | *elongation* | 0.88 | 0.84 | −4.5 / −4.0 | −0.11 / −0.11 | < 0.001 / < 0.001 |
| Drp1 (KO vs. WT) | *area* | 0.92 | 0.98 | +2.3 / −6.1 | 0.01 / −0.04 | 0.85 / 0.22 |
| Drp1 (KO vs. WT) | *elongation* | 0.94 | 0.91 | +14.8 / +15.7 | 0.23 / 0.23 | < 0.001 / < 0.001 |

Three datasets from the study are shown as representative examples. Object counts (U-Net / U-Net++): HCI-010, 26,491 / 29,042 across 346 images; BCM-2147, 9,109 / 9,589 across 181 images; *Drp1*, 1,807 / 1,951 across 63 images. Per-sample *r*, Pearson correlation of image-level mean metric between models. Paired per-object *r*, Pearson correlation on objects matched by centroid proximity (match rate ≥ 94% in all datasets). Δ median %, within-model percent difference of the median (condition − reference). Cliff’s δ thresholds: |δ| < 0.147, negligible; < 0.33, small. Mann–Whitney *U* test, two-sided.

For every metric–dataset combination in which either model detected a non-negligible effect, the two models agreed on direction, effect-size category, and significance. The one sign discrepancy (*Drp1* area) occurs in a null comparison where both |δ| ≤ 0.04 and both *p* > 0.2.
